# Human recombinant erythropoietin improves motor function in rats with spinal cord compression myelopathy

**DOI:** 10.1101/577130

**Authors:** Takahiro Tanaka, Hidetoshi Murata, Ryohei Miyazaki, Tetsuya Yoshizumi, Mitsuru Sato, Makoto Ohtake, Kensuke Tateishi, Phyo Kim, Tetsuya Yamamoto

## Abstract

**OBJECTIVE:** Erythropoietin (EPO) is a clinically available hematopoietic cytokine. The aim of this study was to evaluate the effect of EPO on a rat model of cervical cord compression myelopathy and to explore the possibility of its use as a pharmacological treatment.

**METHODS:** To produce the chronic cervical cord compression model, thin polyurethane sheets were implanted under the C5-C6 laminae of rats and gradually expanded due to water absorption. In this model, motor functions significantly declined from 7 weeks after surgery. Based on the result, EPO administration was started 8 weeks after surgery. Motor function as seen with rotarod performance and grip strength was measured 16 weeks after surgery, and then motor neurons were stained with H-E and NeuN staining, and counted. Apoptotic cell death was assessed with terminal deoxynucleotidyl transferase-mediated deoxyuridine triphosphate-biotin nick end labeling (TUNEL) staining. To assess transfer of EPO into spinal cord tissue, the EPO level in spinal cord tissue was measured with an enzyme-linked immunosorbent assay for each group after subcutaneous injection of EPO.

**RESULTS:** High-dose EPO (5000 IU/kg) administered from 8 weeks after surgery markedly restored and maintained motor function in the Compression groups (P < 0.01). EPO significantly prevented loss of motor neurons in the anterior horn (P < 0.05) and significantly decreased the number of TUNEL-positive apoptotic cells (P < 0.05). The EPO level in spinal cord tissue was significantly higher in the High-dose EPO group than other groups.

**CONCLUSIONS:** EPO improves motor function in rats with progressive chronic compression myelopathy. EPO protects anterior horn motor neurons and inhibits neuronal cell apoptosis in spinal cord compression. The neuroprotective effects can be produced through transfer of EPO into spinal cord tissue. These findings suggest that EPO has high potential as a treatment for developing compression myelopathy.

## Introduction

As the population ages, degenerative changes in the cervical spine progress. The spinal canal gradually narrows due to cervical spondylosis, disc hernia, and ossification of the posterior longitudinal ligament [1, 2]. This chronic compression of the cervical spinal cord causes cervical myelopathy. The symptoms of chronic compression myelopathy such as motor weakness, sensory disturbances, decreased fine motor coordination, and spastic gait gradually progress over time. The main pathogenesis is presumed to be local compression and spinal cord ischemia at the compressed segment [3]. At this time, surgical decompression is often performed to treat cervical compression myelopathy [4–6]. However, no optimal medical treatment is available to improve the neurological status in patients with worsening compression myelopathy.

To elucidate the biological mechanism of chronic compression myelopathy and develop a treatment strategy for it, a co-author, Kim, established a novel experimental model of chronic cervical cord compression [7]. This model is created by inserting a sheet of water-absorbing urethane-compound polymer under the laminae of rats. This model induces delayed motor dysfunction and reproduces the characteristic course of clinical delayed cervical myelopathy. Using this model, we have previously demonstrated that pharmacological agents such as *Limaprost alfadex,* prostaglandin E1 derivative, and *Cilostazol*, a selective type III phosphodiesterase inhibitor, prevent the onset of cervical compression myelopathy [8, 9]. However, functional recovery from developing compression myelopathy has not been elucidated in those studies.

We recently confirmed that granulocyte colony–stimulating factor (G-CSF) improves motor function in the progressive phase of compression myelopathy and preserves anterior horn motor neurons in the rat chronic spinal cord compression model [10]. However, in healthy people, G-CSF causes marked leukocytosis, which commonly results in fever, arthralgia, and rarely, thromboembolism and splenomegaly [11]. Erythropoietin (EPO) is a physiological hematopoietic cytokine like G-CSF. EPO is a 30.4-kDa glycoprotein secreted from the kidney that stimulates red blood cell (RBC) production (erythropoiesis) after binding to the EPO receptor in the bone marrow [12]. EPO is commonly used in anemic patients undergoing chronic hemodialysis or suffering from cancer and undergoing chemotherapy [13, 14]. EPO is also used for preoperative autologous blood donation in hematologically healthy individuals [15]. Therefore, EPO can often be used safely, even in elderly patients or those with critical disease.

In addition, EPO has multifunctional tissue-protective effects, including anti-apoptotic, anti-inflammatory, anti-oxidative, and angiogenic effects [16–18]. During the last two decades, quite a few reports have described its neuroprotective effects in cerebral infarction, brain contusion, and acute spinal cord injury (SCI) in laboratory investigations [19–23]. Those papers reported its effect of neuroprotection, angiogenesis, and anti-apoptosis in the brain and spinal cord [18, 24]. Recently, recombinant human EPO (rhEPO) was preliminarily used in a randomized clinical trial of acute SCI, and results indicated the possibility of treating acute SCI with EPO [25].

However, no reports have shown the neuroprotective effect of EPO for compression myelopathy in experimental or clinical studies.

Here, we investigated the neuroprotective effects of EPO for chronic cervical compression myelopathy using our established rat model of spinal cord compression [7].

## Materials and methods

### Animal maintenance

This study was approved by the Institutional Animal Care and Use Committee of Yokohama City University School of Medicine (IRB: F-A-15-022). Male Wistar rats (12 weeks old, weight 250-300 g; Japan SLC Inc., Hamamatsu, Japan) were housed in cages for 3 weeks before surgery for adaptation to the environment. All rats were trained to exercise on the rotarod device and to undergo forepaw grip strength measurement for 2 weeks before surgery. Throughout this experimental period, the rats had free access to water and food. Body weight was recorded every week during this study.

#### Surgical procedure to create the chronic compression model

The detailed surgical procedure to create the chronic cervical compression model has been described [7]. Under general anesthesia with 2% isoflurane, a midline incision was made in the nuchal area, and the C3-Th1 laminae were exposed. A sheet of expandable urethane compound polymer (size 2 × 6 × 0.7 mm; Aquaprene C^®^, Sanyo Chemical Industries, Ltd., Tokyo, Japan) was inserted into the sublaminar space of C5-C6 (Fig 1A, B). This sheet gradually expands to 230% of the original volume over 48-72 hours by absorbing water in the tissue. In this model, the decline in motor function is delayed, with a latency period after compression introduction, and then gradual progression, whereas no acute damage suggestive of SCI is observed. This model reproduces the characteristic course and features of clinical cervical spondylotic myelopathy [17].

**Fig 1.**
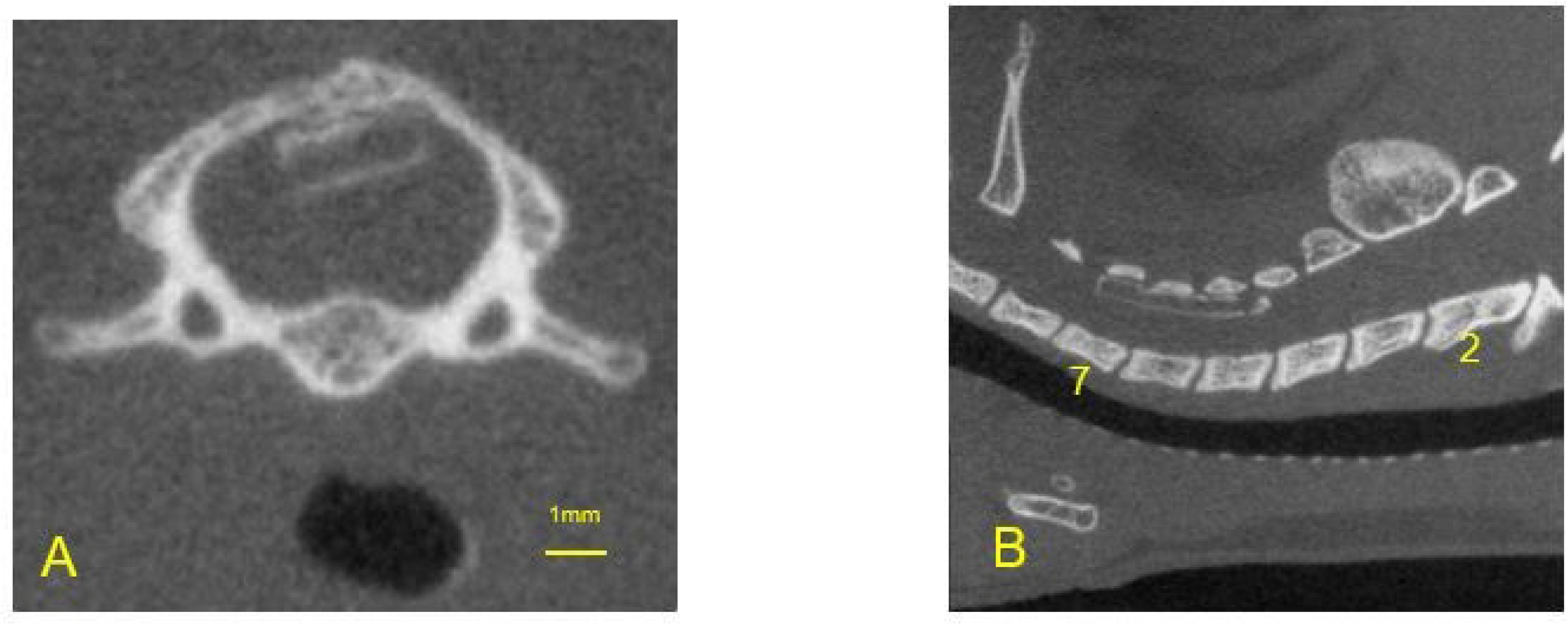
The Chronic Compression Model. A: Computed tomography (CT) axial view at the C5 level, 0.5 mm above the intervertebral foramen. B: CT sagittal view of the cervical spine. Aquaprene^®^ (expandable urethane compound sheet, size 2 × 6 × 0.7 mm) was inserted under the C5-C6 laminae.

### Experimental design

#### Preliminary experiment

As a preliminary experiment, we confirmed the course of motor function decline in this model to determine when to administer EPO in the treatment experiment.

Briefly, 40 rats were allocated to two groups; Sham operation group (n = 15) and Compression group (n = 25). In the Sham group, rats underwent a sham operation; the polymer sheet was placed under the laminae and removed immediately. In the Compression group, this polymer sheet was left in place and continued to compress the spinal cord chronically (Fig 1A, B). The motor functions were evaluated once a week from 1 week before surgery to 26 weeks after surgery.

#### Treatment experiment (Fig 2)

In the treatment experiment, 48 rats were allocated to four groups; Sham group (sham operation + normal saline [NS]; n = 12), Vehicle group (compression + NS; n = 12), Low-dose EPO group (compression + EPO low dose; n = 12), and High-dose EPO group (compression + EPO high dose; n = 12). From the results of the preliminary experiments, the motor function was significantly decreased 8 weeks after surgery. Therefore, administration of rhEPO or NS was started from 8 weeks after surgery and lasted until 16 weeks; the frequency of administration was twice a week. In the Sham group, rats underwent the sham operation and received administration of NS subcutaneously. In the Vehicle group, rats underwent polymer sheet implantation and received administration of NS subcutaneously. In the Low-dose EPO group, cervical compression model rats received rhEPO 500 IU/kg/day (rhEPO; kindly provided by Chugai Pharmaceutical Co., Ltd., Tokyo, Japan) subcutaneously. In the High-dose EPO group, cervical compression model rats received administration of rhEPO 5000 IU/kg/day subcutaneously. The motor functions were also evaluated once a week from 1 week before surgery to 16 weeks after surgery. Histological assessment of the anterior horn was evaluated at 16 weeks after surgery (Fig 2A).

**Fig 2.**
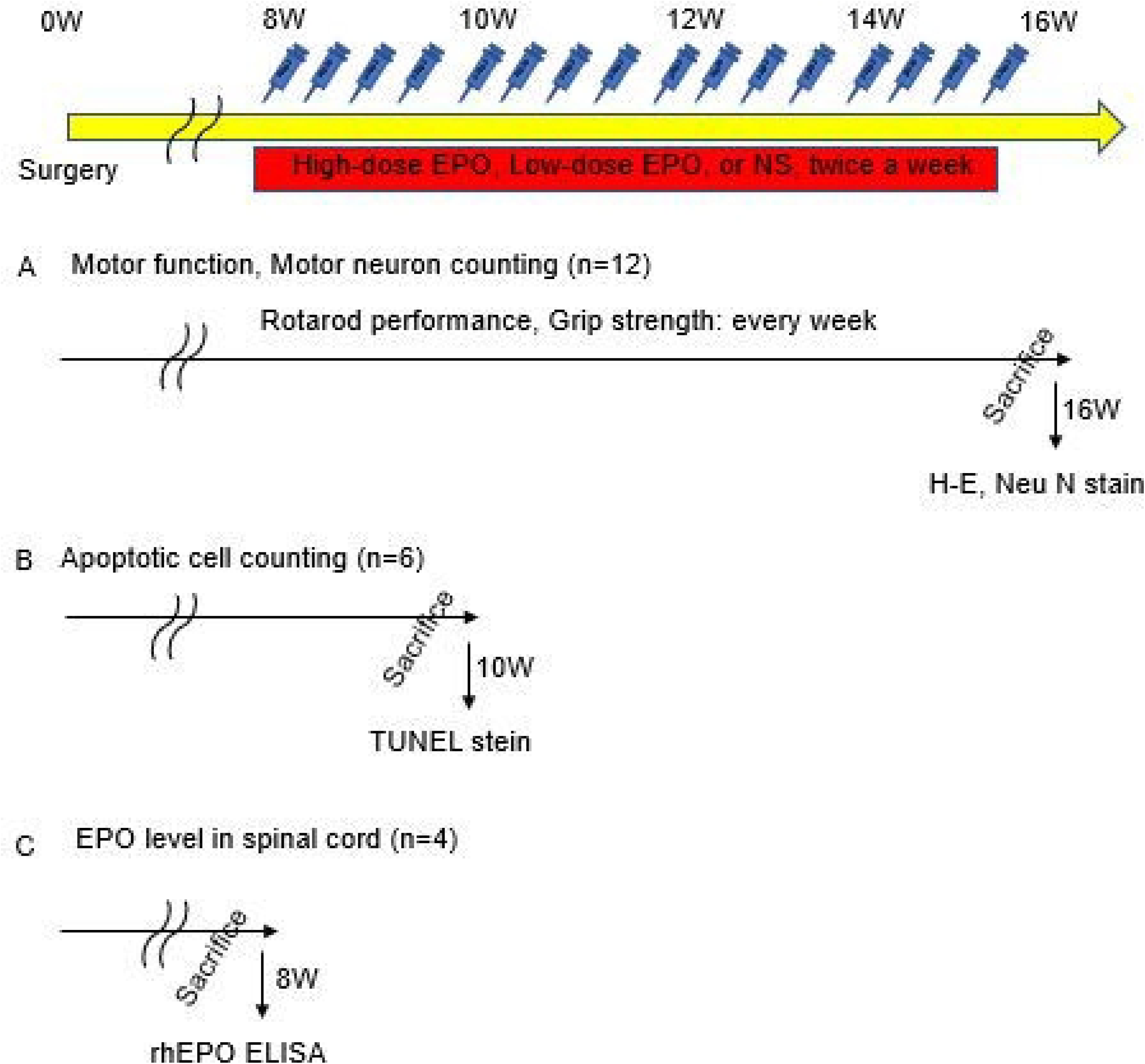
Treatment Experiment: Experimental Design. Administration of low-dose and high-dose EPO and normal saline was started twice a week at 8 weeks postoperatively. **A:** Forty-eight rats were divided into four groups (Sham, Vehicle, High-dose EPO, Low-dose EPO). The motor functions of rotarod performance and grip strength were evaluated once a week before surgery to 16 weeks after surgery. Every rat was sacrificed, and histological analysis was performed (H-E staining and NeuN staining). **B:** Another 18 rats were divided into three groups (Sham, Vehicle, High-dose EPO). Treatment was done from 8 weeks to 10 weeks after surgery, and all rats were sacrificed at 10 weeks after surgery. Apoptotic cells were evaluated with TUNEL staining at 10 weeks after surgery. **C:** Another 12 rats were divided into three groups (Vehicle, High-dose EPO, Low-dose EPO). Each single treatment was done 8 weeks after surgery. All rats were sacrificed 12 hours after injection, and the EPO level in the spinal cord was measured using a rhEPO enzyme-linked immunosorbent assay (ELISA).

#### Motor function analysis

##### Rotarod performance

Rotarod performance was assessed by using the rotarod device (ENV-557, Med Associates Inc., St. Albans, VT). Based on our previous research, a moderate rotation speed of 10 rpm was set [7–10]. All rats could walk on the rotarod for more than 300 seconds before surgery. Therefore, 300 seconds was set as the cut-off. Three trials in each session were performed for all rats. We recorded the longest duration time of the three trials.

##### Forelimb grip strength

Forelimb grip strength was assessed by using a digital force meter (MK-380CM/F, Muromachi Kikai, Tokyo, Japan). We assessed grip strength according to the methods of Meyer et al [26]. The animals were evaluated before surgery and once a week after surgery. All rats also performed three trials in each session, and the maximum score (in newtons: N) was used for data analysis.

#### Histological analysis

##### Hematoxylin and eosin (H-E) staining

At 16 weeks after surgery, transcardial perfusion was performed with 4% paraformaldehyde in phosphate-buffered saline (PBS) in all rats. The spinal cord segment at C5-6 was removed en bloc and placed in 4% paraformaldehyde solution for 3 days. After this process, these C5-6 segments were embedded in paraffin and sectioned at a slice thickness of 5 µm and a gap interval of 5 µm over 1000 µm length, according to stereological considerations of motor neurons [7–10]. One hundred specimens of all rats were stained with H-E. Motor neurons have large nuclei and well-developed, densely stained Nissl bodies in the cytoplasm. The characteristic large nucleolus has a uniform diameter of approximately 5 µm [27, 28]. In H-E-stained sections, we regarded such cells as motor neurons. Motor neurons on both sides of the anterior horn gray matter were counted.

##### NeuN staining

NeuN protein appears in neuron-specific nuclei. The nucleus of the motor neuron is more clearly detected with NeuN staining compared with H-E staining.

In the treatment experiment, 10 specimens (thickness = 5 µm, gap interval = 50 µm) of five rats in all groups were stained with immunohistochemistry. Both sides of the anterior horn were also evaluated with NeuN staining following application of the chromogen diaminobenzidine (Dako North America, Santa Clala, CA, USA, 1:100) using the labeled streptavidin biotin technique [29]. Rabbit anti-NeuN (EMD Millipore Corporation, Burlington, MA, USA, 1:100) was used as the primary antibody. NeuN-positive cells on both sides of the anterior horn gray matter were counted.

##### Terminal deoxynucleotidyl transferase-mediated deoxyuridine triphosphate-biotin nick end labeling (TUNEL) staining

Apoptotic cell death was investigated 10 weeks after surgery. Another 18 rats (Sham group; n = 6, Vehicle group; n = 6, high-dose EPO group; n = 6) were perfused transcardially with 4% paraformaldehyde in PBS (Fig 2B). The C5-6 segment of the spinal cord was embedded in optimal cutting temperature compound and frozen in liquid nitrogen. Three sections from C5-6 segments (thickness = 20 µm, gap interval = 50 µm) were cut in a cryostat and stained with the In Situ Cell Death Detection Kit, POD (Roche, Basel, Switzerland) according to the manufacturer’s recommendations. Nuclei were counterstained with 4’,6-diamidino-2-phenylindole (DAPI, 1:5000 in PBS) (Molecular Probes, Eugene, OR). The TUNEL stain signal was observed under an FV300 confocal microscope (Olympus Optical Company, Ltd., Tokyo, Japan). TUNEL- and DAPI-positive cells were counted, and the ratios of apoptotic cells to total nuclei were evaluated in each group.

### Hematological assessment

Another 12 rats were divided into three groups. All these rats underwent the operation to place the polymer sheet under the C5-6 laminae and were treated with NS or rhEPO twice a week from 8 weeks after surgery. The Vehicle group, Low-dose EPO group, and High-dose EPO group were examined. All rats were subjected to inhalation anesthesia with 2% isoflurane. Blood samples (0.5 ml/body) were collected by venipuncture from the tail vein at 2, 4, and 6 weeks after the first EPO administration. Blood samples were collected into blood collection tubes with EDTA 2K (BD Microtainer, Japan Becton, Dickinson and Company, Tokyo, Japan) immediately. RBC, hemoglobin (Hb), and hematocrit (Ht) values were assessed with an automated hematology analyzer (XE 2100, Sysmex, Hyogo, Japan).

### The rhEPO level in spinal cord tissue

To assess whether subcutaneously injected EPO was transferred to the spinal cord, we measured EPO levels in the spinal cord with an enzyme-linked immunosorbent assay. Another 12 rats were divided into three groups; Vehicle group, Low-dose EPO group, and High-dose EPO group. All these rats underwent the operation in which the polymer sheet remained under the C5-6 laminae. They received NS or rhEPO 8 weeks after surgery. Twelve hours after subcutaneous injection of NS or rhEPO, all rats were sacrificed under anesthesia, and blood was completely removed by transcardial perfusion with PBS to exclude rhEPO from blood (Fig 2C). The spinal cord segment at the C5-6 level was removed en bloc. These tissues were homogenized in IP buffer with an ultra Turrax homogenizer and centrifuged at 12000 rpm at 4°C for 5 min. Supernatants were removed and analyzed to determine the levels of rhEPO in the spinal cord tissue. The total protein of the spinal cord tissue was determined using bovine serum albumin as a standard. The rhEPO concentration in spinal cord tissue was measured with a rhEPO enzyme-linked immunosorbent assay kit (R&D Systems Europe, Abingdon, UK) according to the manufacturer’s instructions. The concentration was described as the rhEPO level per 1 g tissue (mIU/g) and tissue dose % of injected dose (%ID).

### Statistical Analysis

GraphPad Prism 6 software for Windows (GraphPad Software Inc., La Jolla, CA, USA) was used for statistical analysis. Data are expressed as the means ± standard error of the mean. The duration of walking on the rotarod, forelimb grip strength, and hematological data were analyzed using two-way repeated-measures analysis of variance. The number of anterior horn motor neurons with H-E, NeuN, and TUNEL staining and the rhEPO level in spinal cord were tested using one-way analysis of variance. P values <0.05 were regarded as significant.

## Results

### Motor function

#### Preliminary experiment

Rotarod performance declined gradually with a latency period of 4 weeks in the Compression group. At 7 weeks after surgery, the walking duration significantly decreased in the Compression group compared to the Sham group (P < 0.001). In the Compression group, the duration declined gradually and reached a plateau after 16 weeks (Fig 3A).

**Fig 3.**
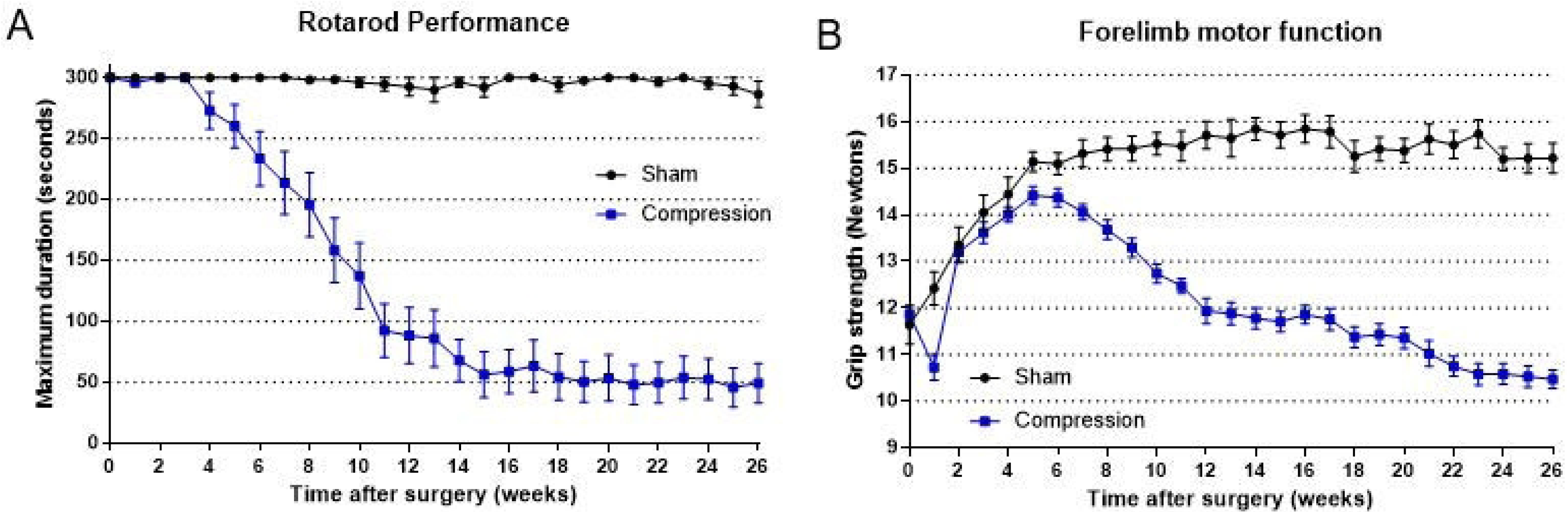
Preliminary experiment. A: Time course of rotarod performance measured by walking time on a rotarod (cut-off 300 seconds). In the Compression group, the walking time gradually started to decline from 4 weeks, and showed a significant decrease at 7 weeks after surgery (P < 0.001). The performance reached a plateau with a low duration of about 50 seconds after 15 weeks. B: Time course of grip strength. In the Compression group, grip strength decreased at 1 week after surgery, but gradually increased as body weight increased. However, grip strength gradually declined from 6 weeks, and showed a significant difference at 7 weeks after surgery (P < 0.0001) After that, the strength continued to decrease gradually, reaching approximately 10.5 N at 26 weeks postoperatively.

Forelimb grip strength increased until 5 weeks after surgery and started to decrease from 6 weeks in the Compression group. The strength gradually declined and significantly decreased after 7 weeks compared with the Sham group (P < 0.001) (Fig 3B).

Based on these results, we decided to administer EPO beginning 8 weeks after surgery as a treatment experiment.

#### Treatment experiment

The rotarod performance of the Compression groups (Vehicle, Low-dose EPO, High-dose EPO groups) declined gradually from 5 weeks after surgery, and a significant decrease was seen from 7 weeks compared with the Sham group (P < 0.005) as in the preliminary experiments (Fig 3A).

After EPO administration beginning 8 weeks after surgery, rotarod performance started to improve in the treatment groups (Low-dose EPO, High-dose EPO group). Especially in the High-dose EPO group, rotarod performance significantly improved compared with the other Compression groups (Vehicle, Low-dose EPO groups), although the performance gradually declined from 13 weeks. The effects of EPO continued for 5 weeks after EPO administration (P < 0.01). Furthermore, the High-dose EPO group improved to the level at which no significant difference in motor function was seen between the Sham and High-dose EPO groups at 9 weeks after surgery. The Low-dose EPO group showed slightly improved rotarod performance, but did not show significant improvement compared with the Vehicle group (Fig 4A).

**Fig 4.**
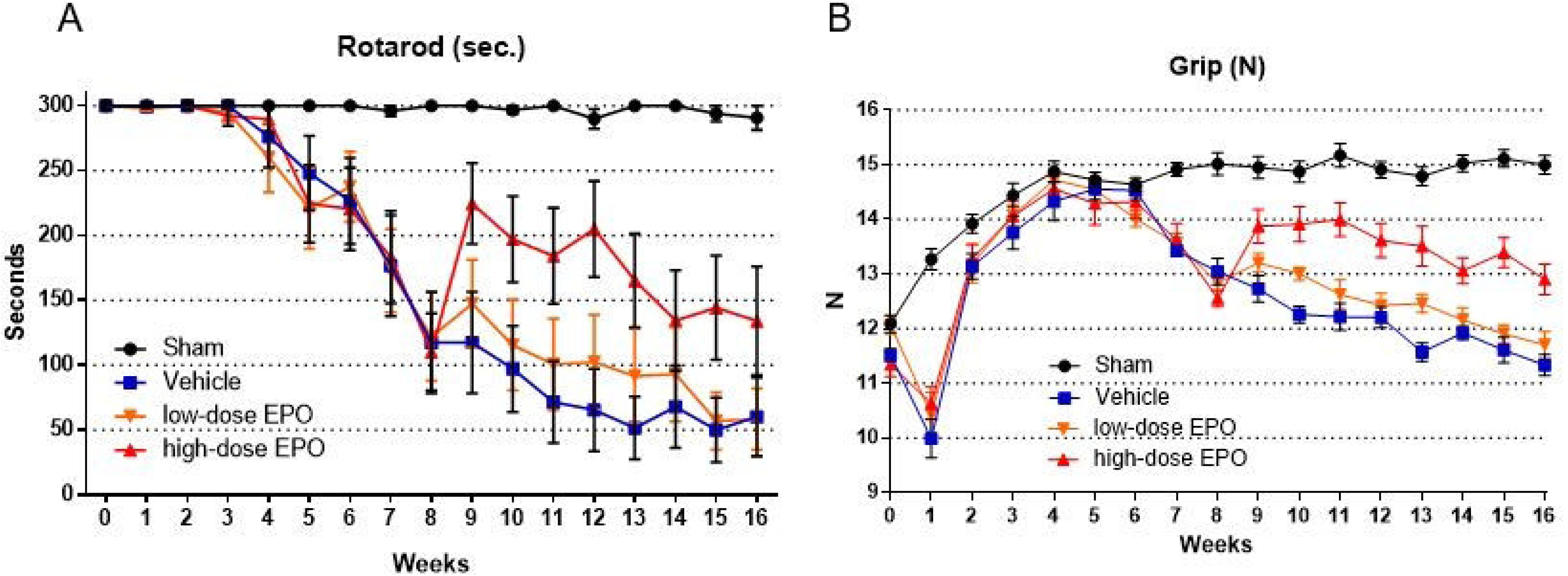
Treatment experiment: Motor function. A: Time course of rotarod performance measured by walking time on a rotarod (cut-off 300 seconds). In the Compression models (Vehicle, Low-dose EPO, and High-dose EPO groups), rotarod performance gradually declined from 3 weeks after surgery, and showed a significant difference at 7 weeks after surgery. After administration of EPO from 8 weeks, rotarod performance improved in the EPO groups. Especially in the High-dose EPO group, performance markedly improved. This effect was maintained with a significant difference by week 13 after surgery (P < 0.01). In the Low-dose EPO group, slight improvement in rotarod performance was observed, but it did not reach statistical significance compared with the Vehicle group. B: Time course of grip strength. In the Compression groups, the strength started to decline from 6 weeks after surgery, and a significant decline was observed at 7 weeks. EPO was administered at 8 weeks, and grip strength improved, especially in the High-dose EPO group. Significant improvement was seen from 9 weeks in the High-dose EPO group (P < 0.0001) and continued up to 16 weeks after surgery. In the Low-dose EPO group, grip strength slightly improved, but no significant difference was found compared with the Vehicle group.

The forelimb grip strength of the Compression groups decreased at 1 week after surgery but started to recover gradually from 2 weeks after surgery. The strength of the Compression group showed an improvement course equal to that of the Sham group from 2 weeks after surgery and then started to decrease gradually from 7 weeks; at this time, the strength was significantly decreased compared with the Sham group (P < 0.001) (Fig 4B).

After EPO administration at 8 weeks after surgery, grip strength started to improve in the treatment groups (Low-dose EPO, High-dose EPO groups).

In the High-dose EPO group, the strength significantly improved compared with the other Compression groups (Vehicle, Low-dose EPO groups) (P < 0.0001). Its effects continued throughout the period of EPO administration (9 to 16 weeks after surgery), although the strength gradually decreased from 4 weeks after EPO administration.

In contrast, the Low-dose EPO group showed a slight improvement in strength, but did not show significant improvement compared with the Vehicle group (Fig 4B).

### Histopathological analysis

#### H-E staining

At 16 weeks after surgery, loss of anterior horn motor neurons and vacuolar degeneration in the spinal cord were observed in H-E-stained sections from the Compression groups (Vehicle, Low-dose EPO, and High-dose EPO group) (Fig 5A). The numbers of motor neurons were 1834.7 ± 115.4 (Sham group), 1421.6 ± 50.1 (Vehicle group), 1484.7 ± 74.2 (Low-dose EPO group), and 1640.0 ± 66.9 (High-dose EPO group). The number of motor neurons on both sides of the anterior horn was significantly decreased in every Compression group compared to the non-compression Sham group (P < 0.0001). In the High-dose EPO group, however, the motor neurons were significantly preserved compared with the other Compression groups (Vehicle and Low-dose EPO group; P < 0.0001, P < 0.0005) (Fig 5B).

**Fig 5.**
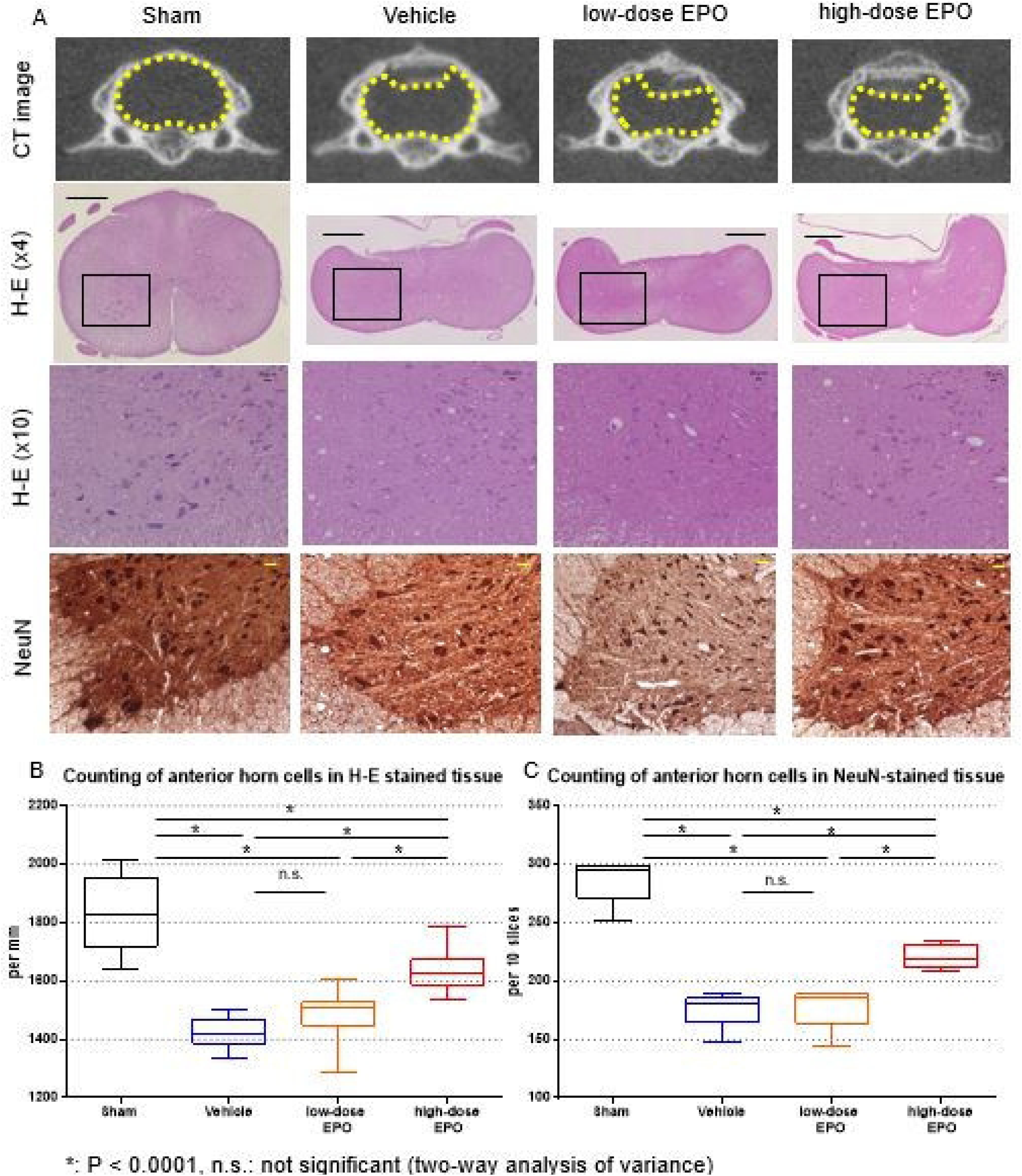
Treatment experiment: Anterior motor neurons. A: Top panels: CT axial view in C5. In the Compression groups (Vehicle, Low-dose EPO, and High-dose EPO), the spinal cord was compressed by Aquaprene^®^ (expandable urethane compound sheet, size 2 × 6 × 0.7 mm). The yellow dotted figure shows the outline of the spinal cord. Second panels: The spinal cord at the C5 level was sliced into 5-μm thick sections at 16 weeks after surgery. Hematoxylin and eosin staining of cross sections of spinal cord is shown (original magnification ×4, scale bar = 100 µm). In the Compression groups, the spinal cord was flattened. Black box shows the region of the anterior horn. Third panels: The black box in the second panel was magnified (×10, scale bar 20 µm). Cells with large nuclei and well-developed, densely stained Nissl bodies in the cytoplasm indicate motor neurons. In the Vehicle and Low-dose EPO groups, motor neurons decreased, and vacuolar degeneration was obvious. In the High-dose EPO group, motor neurons were preserved, although vacuolar degeneration was present. Bottom panels: NeuN staining of the anterior horn (×10, scale bar 20 µm). The nuclei of motor neurons are clearly detected with NeuN staining compared with H-E staining. Motor neurons decreased in the Vehicle and Low-dose EPO groups, but were preserved in the High-dose EPO group. B: Counting of anterior horn cells in H-E-stained tissue. The number of cells with large nuclei in the anterior horn was counted in every group. The number was significantly decreased in the Compression groups compared with the Sham group (*P < 0.0001). However, in the High-dose EPO group, the number was significantly preserved compared with the other two Compression groups (Vehicle and Low-dose EPO group) C: Counting of anterior horn cells in NeuN-stained tissue. NeuN-positive cells were significantly decreased in the Compression groups compared with the Sham group (*P < 0.0001). However, in the High-dose EPO group, the number was significantly preserved compared with the other two Compression groups (*P < 0.0001). The tendency in the cell count was similar to that with H-E staining.

#### Cell counting of NeuN-positive cells

The number of NeuN-positive cells in 10 slices of each group was 286.8 ± 17.6 (Sham group), 176.0 ± 14.3 (Vehicle group), 178.0 ± 17.1 (Low-dose EPO group), and 220.4 ± 9.4 (High-dose EPO group). NeuN-positive cells in each Compression group decreased compared with the Sham group, but the number in the High-dose EPO group was significantly preserved compared with the Vehicle and Low-dose EPO groups. This tendency was similar to that of the number of motor neurons in H-E-stained sections (Fig 5B, 5C).

#### TUNEL staining

TUNEL-positive cells were significantly increased in the Vehicle group compared with the other two groups (Sham and High-dose EPO groups) (P < 0.0001) (Fig 6A, 6B). We found no significant difference between the Sham and High-dose EPO groups. The ratios of TUNEL-positive cells to DAPI-positive cells (%) were 1.72 ± 0.59% (Sham group), 35.01 ± 9.17% (Vehicle group), and 5.66 ± 2.27% (High-dose EPO group). The ratio in the Vehicle group was significantly higher than that in the other two groups (P < 0.0001), and we found no significant difference between the Sham and High-dose EPO groups (Fig 6A, 6C).

**Fig 6.**
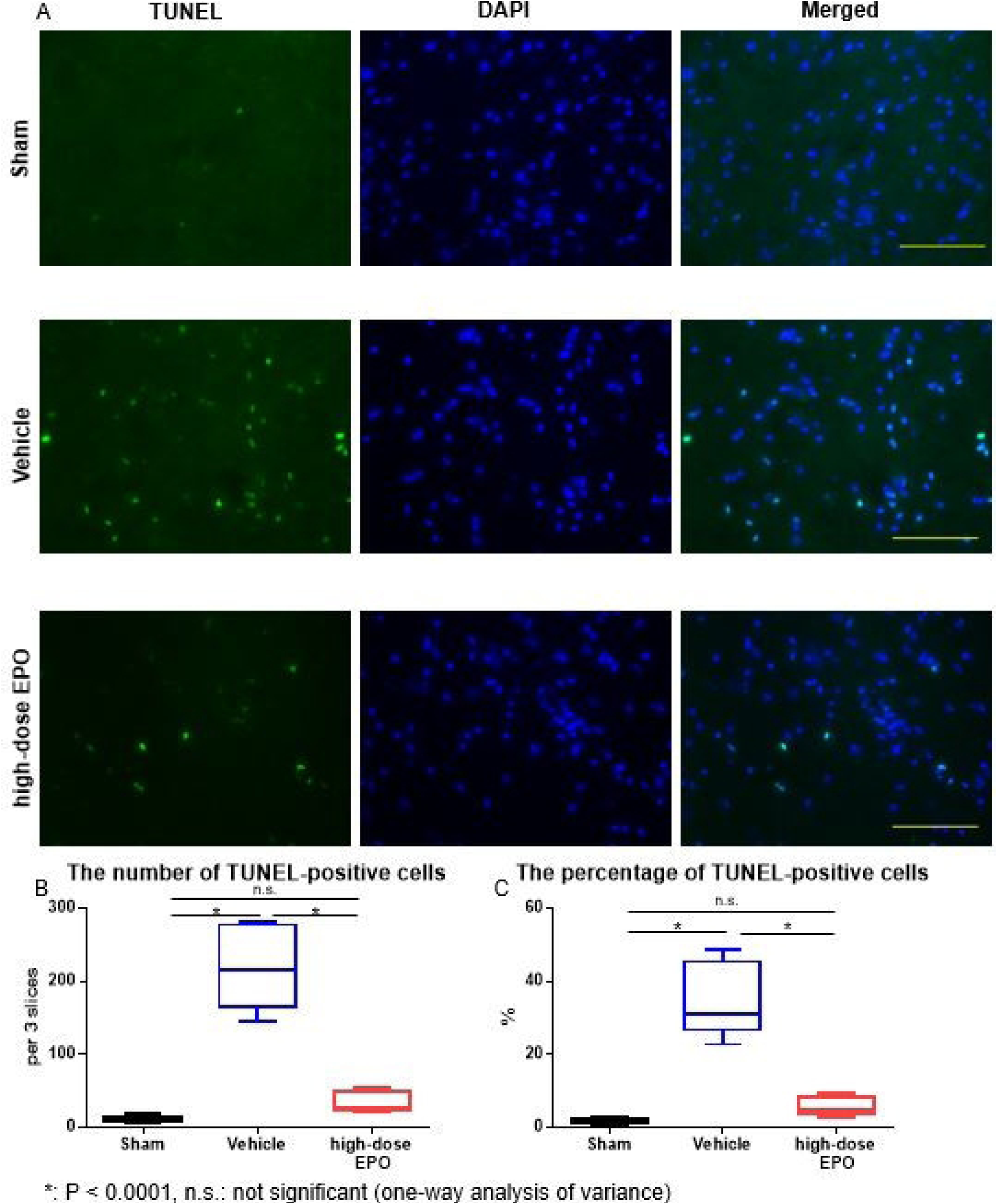
Treatment experiment: TUNEL staining. A: TUNEL staining was performed to detect apoptotic cells at 10 weeks after surgery. DAPI/TUNEL double staining is shown in each group (DAPI staining, TUNEL staining, DAPI/TUNEL staining, Bar = 100 µm). The Vehicle group showed the highest number of TUNEL-positive cells. B: The number of TUNEL-positive cells was counted in each group. The number of TUNEL-positive cells in the Vehicle group was significantly higher than in the other two groups (*P < 0.0001), with no significant difference between the Sham group and High-dose EPO group. C: The percentage of TUNEL-positive cells in each group. The percentage in the Vehicle group was significantly higher than in the other two groups (*P < 0.0001).

### Hematological data

After administration of EPO, the RBC, Hb, and Ht values increased immediately in the EPO-administered groups (Low-dose and High-dose EPO groups) (P < 0.0001). The trend in RBC and Hb values showed a similar increasing tendency after EPO administration (Fig 7A, B). Eventually, the RBC and Hb values increased to approximately 1.2 and 1.4 times in the Low-dose and High-dose EPO groups, respectively, compared to the baseline value (Vehicle group). The values were significantly higher in both EPO-administered groups than the Vehicle group until 6 weeks after administration (P < 0.0001) (Fig 7A, B).

**Fig 7.**
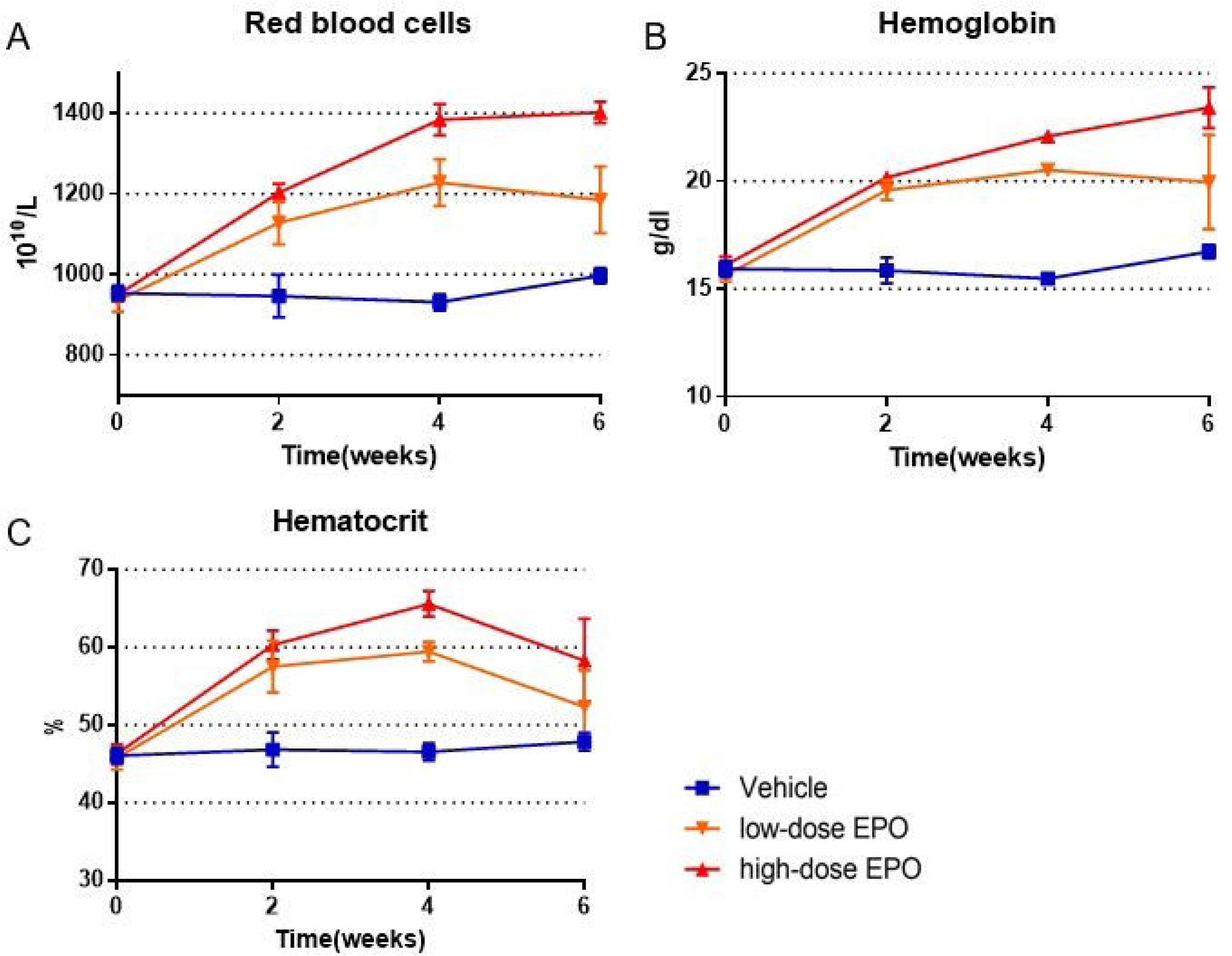
Treatment experiment: Hematological data. A: Time course of red blood cells (RBCs). RBCs increased immediately in the Low-dose and High-dose EPO groups (P < 0.0001). From 4 weeks after administration, we found a significant difference between the Low-dose and High-dose EPO groups (P < 0.0001). Eventually, RBCs increased up to approximately 1.2 and 1.4 times in the Low-dose and High-dose EPO groups, respectively, compared with the Vehicle group. B: Time course of hemoglobin (Hb). The Hb value increased immediately in the Low-dose and High-dose EPO groups (P < 0.0001). The time course was similar to that of RBCs. Eventually, the Hb value increased up to approximately 1.2 and 1.4 times in the Low-dose and High-dose EPO groups, respectively, compared with the Vehicle group. C: Time course of hematocrit (Ht). The Ht value increased immediately in the Low-dose and High-dose EPO groups (P < 0.0001). The Ht value was the highest at 4 weeks, and then peaked. The maximum Ht value was approximately 1.3 and 1.4 times in the Low-dose and High-dose EPO groups, respectively, compared with the Vehicle group.

The Ht value in the EPO-administered groups was the highest at 4 weeks and increased to approximately 1.3 and 1.4 times in the Low-dose and High-dose EPO groups, respectively, compared to the baseline value (P < 0.0001). At 6 weeks after administration of EPO, the Ht value of the EPO-administered groups started to peak. The Ht value of the High-dose EPO group was significantly higher than that of the other two groups (P = 0.005) at 6 weeks (Fig 7C).

### rhEPO level in spinal cord tissue

The rhEPO level in the spinal cord 12 hours after subcutaneous injection of rhEPO was less than 0.10 mIU/g in the Vehicle group, 1.07 ± 0.46 mIU/g in the Low-dose EPO group, and 8.67 ± 2.33 mIU/g in the High-dose EPO group. The rhEPO level was remarkably higher in the High-dose EPO group than the other two groups (P < 0.0001). In the Low-dose EPO group, the rhEPO level was slightly increased, but that of the Low-dose EPO group did not show a significant difference compared with the Vehicle group (Fig 8A).

**Fig 8.**
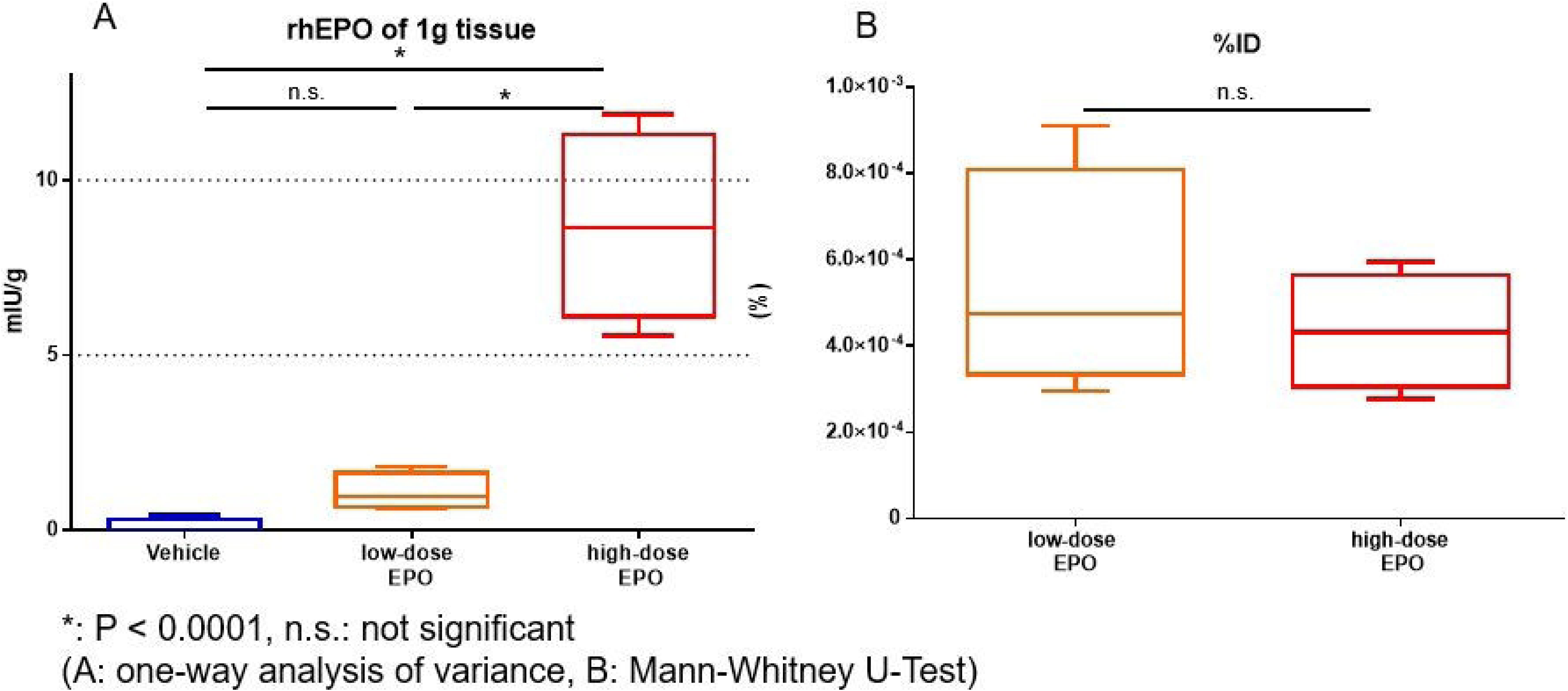
Treatment experiment: ELISA of recombinant human EPO (rhEPO) A: The amount of rhEPO per 1 g spinal cord tissue. The rhEPO level was significantly higher in the High-dose EPO group compared to the other two groups (*P < 0.0001). B: The tissue rhEPO level for injected dose (%ID) in the Low-dose and High-dose EPO groups. No significant difference was found in the %ID between the two groups.

The tissue % ID was 4.4 ± 1.2 (10^−4^%) in the High-dose EPO group and 5.4 ± 2.3 (10^−4^%) in the Low-dose EPO group. We found no significant difference between the two groups (Fig 8B). This result shows that the rhEPO level in the spinal cord was dose dependent.

## Discussion

The present study demonstrated that EPO improved motor functions and preserved motor neurons, even in developing myelopathy due to spinal cord compression. Furthermore, EPO was transferred into spinal cord tissue following subcutaneous EPO administration.

Some studies reported that EPO improves motor function in an experimental acute SCI model [21, 22, 30, 31]. EPO and the EPO receptor (EPO-R) are highly expressed in both the central and peripheral nervous systems [32]. The roles of EPO in these areas are in neuroprotection, angiogenesis, anti-apoptosis, and anti-inflammation [18, 24, 33]. Clinically, a preliminary randomized comparative trial was performed in patients with acute SCI. In this trial, the effect of EPO treatment was compared with high-dose methylprednisolone treatment. EPO had higher efficacy and fewer side effects than methylprednisolone, indicating a therapeutic effect for acute SCI patients [25].

In contrast, the neuroprotective effect of EPO for compression myelopathy remains unknown. We previously demonstrated that blood flow in the compressed segment is markedly reduced, indicating the presence of local spinal cord ischemia in the chronic compression myelopathy model [32]. Consistent with our previous studies [9, 10], this study also demonstrated that chronic spinal cord compression induces apoptotic cell death. In hypoxic stress conditions, endogenous EPO is produced in response to low oxygen partial pressure and protects neurons [34]. Importantly, cell apoptosis induced by spinal cord compression is inhibited by high-dose EPO administration, indicating anti-apoptosis and anti-inflammatory effects of EPO [18, 24, 33]. Additionally, a rapid increase in RBC, Hb, and Ht values following EPO administration may improve the local oxygen supply and restore motor function (Fig 7). Liem et al. reported that blood transfusion for anemia improves cerebral oxygenation in newborn infants [35]. Although we could not directly evaluate local oxygen pressure, we speculate that improvement in cervical myelopathy is due to anti-apoptotic effects of EPO and improvement in local ischemia in the spinal cord with an increased oxygen supply.

In the current study, both high-dose and low-dose EPO increased hematopoietic values including RBC, Hb, and Ht. However, functional recovery was observed with high-dose EPO treatment in particular. High-dose EPO may have passed through the blood-spinal cord barrier. EPO is a high-molecular weight glycoprotein (30.4 kDa) [12]. In classic papers, the blood–brain barrier (BBB) was considered to be impermeable to large glycosylated molecules like EPO [36]. However, some recent studies have reported that EPO can pass through the BBB due to a high concentration and after BBB disruption such as that which follows brain and spinal cord contusion [37–39]. EPO can cross the BBB at 450 IU/kg or more in rats [37] and crosses the BBB in a dose-dependent manner in a rat brain contusion model [40]. In the current study, in fact, high-dose EPO was predominantly transferred into the spinal cord tissue 12 hours after EPO subcutaneous administration, probably resulting from passing through the blood-spinal cord barrier. Transfer of EPO into spinal cord tissue was dose dependent (Fig 8A), and the transfer activity was almost the same between the Low-dose and High-dose EPO groups (Fig 8B). This finding demonstrates that the higher the dose of EPO that was administered, the more EPO can transfer into spinal cord tissue. This result indicates that EPO directly affected the spinal cord to provide neuronal protection as well as indirectly affected the cord by increasing RBC, Hb, and Ht values.

The dosage of EPO (500 IU/kg or 5000 IU/kg) in this study was decided based on previous reports in acute or subacute SCI with no side effects including hematological complications [21, 41–43].

EPO has been used in clinical practice for a long time, and knowledge of the hematopoietic effect, clinical safety, and side effects of EPO has accumulated. The possible side effects of EPO in humans include hypertension, coagulation disorders, and polycythemia. [44] However, no adverse effects occurred in brain injury patients treated with 10000 IU/kg for 7 consecutive days [45]. In a recent preliminary randomized comparative trial (EPO versus methylprednisolone) for human acute SCI, EPO (500 IU/kg) had a predominant effect and no adverse effects compared with high-dose methylprednisolone. Based on these data, EPO may be a clinically acceptable agent for progressive compression myelopathy as well as a hematopoietic cytokine.

Polycythemia vera (erythemia) is defined as a Hb value more than 18.5 in males and 16.5 in females by WHO guidelines [46]. In practical clinical use, EPO should be used while monitoring of RBC, Hb, and Ht values, especially in hematologically healthy people. Administration of EPO is indicated for patients with anemia and those waiting for surgery and expecting preoperative hematopoietic effects.

The effect of EPO treatment gradually declined at 4 weeks after EPO administration in this rat model of compression myelopathy, although the group given high-dose EPO was finally superior to the group given NS in terms of motor functions. Therefore, the best treatment period may be limited to several weeks after EPO administration, and surgical decompression may be considered during that period.

Certainly, continuous administration of EPO to patients with simple cervical spondylosis over a long period seems unrealistic considering the side effects and high costs. Practical clinical use of EPO may occur for a limited period, especially in patients with worsening symptoms of compression myelopathy who have higher systemic risks such as severe anemia, older age, and diabetes mellitus, and those who live far from a hospital that performs spinal surgery. Furthermore, EPO may be effective against surgical complications such as compression myelopathy due to postoperative epidural hematoma and spinal alignment failure.

The detailed mechanisms of the neuroprotective effect of EPO for compression myelopathy remain to be elucidated. In addition, in this study, the changes in local blood flow and oxygen partial pressure in the spinal cord were not elucidated. However, this study strongly suggests that EPO has potential for treating patients with developing compression myelopathy, and may be worth reconsidering for clinical use to provide both neuroprotective and hematopoietic effects. Further investigations including larger randomized controlled trials with long-term follow-up surveys are required to establish the clinical efficacy of EPO treatment and elucidate therapy-related adverse events.

## Conclusions

EPO improved motor function in rats with developing myelopathy due to chronic spinal cord compression. EPO protected anterior horn motor neurons and decreased neuronal cell apoptosis. The neuroprotective effects were produced following transfer of EPO into the spinal cord tissue. These findings suggest that EPO has high potential as a treatment for developing compression myelopathy.

## Notes

**Conflicts of Interest:** Human Recombinant Erythropoietin was kindly provided by Chugai Pharmaceutical Co., Ltd., Osaka, Japan. We do not receive any research funding from this company.

**Source of Funding:** This work was supported in part by The General Insurance Association of Japan and a Grant-in-Aid for Scientific Research of Japan (17K10903).

